# Changes in anterior cingulate connectivity associated with response to antidepressant treatment for major depressive disorder

**DOI:** 10.64898/2026.10.29.762937

**Authors:** Choi Sze Tracy Lam, Manuel Kuhn, Lauren R. Borchers, Boyu Ren, Sarah E. Woronko, Mohan Li, Ethan M. Zhang, Brian P. Brennan, Gordana Vitaliano, Ty Lees, Mario Bogdanov, Nikolaos P. Daskalakis, Kerry J. Ressler, Peter Zhukovsky, Diego A. Pizzagalli

## Abstract

**Background:** Resting-state functional connectivity (rsFC) involving the anterior cingulate cortex (ACC) and the hippocampal formation has been implicated in major depressive disorder (MDD) and antidepressant response. However, it remains unclear whether connectivity changes in distinct ACC subdivisions and hippocampal formation reflect general antidepressant effects or are specifically associated with clinical improvement.

**Methods:** Forty-one, unmedicated adults with MDD underwent functional MRI before and after 8 weeks of sertraline or bupropion treatment. Seed-based rsFC analyses examined subgenual ACC (sgACC), pregenual ACC (pgACC), dorsal ACC (dACC), and hippocampal formation connectivity. Linear mixed-effects models tested whether connectivity change was associated with treatment response, remission, and dimensional improvement in depressive symptoms.

**Results:** Thirty individuals (73.2%) responded to treatment and 26 (63.4%) achieved remission. Overall, rsFC decreased pre- to post-treatment between (1) dACC and pgACC (but not sgACC and hippocampal formation) and (2) cortical regions primarily involving default mode and frontoparietal networks. When considering treatment response, differences in sgACC and pgACC connectivity with temporal and inferior frontal regions emerged (increased in responders vs. decreased in non-responders). Remission was similarly associated with increased sgACC connectivity with temporal and prefrontal regions, and increased sgACC connectivity with temporal cortical regions related to greater reductions in depressive symptoms.

**Conclusions:** Antidepressant treatment was associated with distinct patterns of ACC connectivity change, with general reductions in dorsal and pregenual connectivity but increases in subgenual and pregenual connectivity associated with favorable clinical outcomes. Findings suggest that changes in sgACC/pgACC connectivity are particularly relevant to antidepressant treatment response.

## INTRODUCTION

Major depressive disorder (MDD) is a highly prevalent psychiatric condition [1], characterized by dysfunction in brain networks spanning affective processing, cognitive control, salience detection, reward processing, and self-referential cognition [2, 3]. Among these circuits, anterior cingulate cortex (ACC) subregions and hippocampal regions play a critical role in the course of depressive episodes [3-6]. In particular, disrupted coordination of the ACC with the default mode and frontoparietal networks may contribute to difficulties in emotion regulation, persistent negative self-referential processing, and reduced behavioral flexibility [2, 7-9]. Antidepressant treatment has been associated with region-specific connectivity changes, including decreased dorsomedial prefrontal and increased hippocampal connectivity following escitalopram [10], and predominantly reduced connectivity across distributed networks among patients achieving remission with medication [11]. Early increase in visual-precuneus-thalamic connectivity has also been observed following both antidepressant and placebo treatment, irrespective of clinical outcome [12]. However, these neural changes are heterogeneous, consistent with substantial variability in antidepressant treatment outcomes, with many patients not responding to one or more depression treatments [13,14]. Understanding the neural correlates of response to depression treatment is critical to advancing efforts to improve clinical outcomes [11,12].

The ACC has been central to efforts to characterize the neural circuitry underlying antidepressant response. Connectivity involving the pregenual (rostral) and subgenual ACC has been associated with symptom improvement, remission, and differential outcomes across antidepressant medications and psychotherapy [15-19]. However, the ACC is not a unitary structure. Cytoarchitectonic, connectional, and functional studies distinguish subgenual (sgACC), pregenual (pgACC), and dorsal ACC (dACC) subregions with partially dissociable roles in affective, visceromotor, salience, and cognitive-control circuits [4, 20-23]. These subdivisions participate in distinct but interconnected circuits, with the sgACC implicated in affective and visceromotor regulation, the pgACC in affective evaluation and integration, and the dACC in cognitive control and salience-related processing. Consistent with this functional differentiation, neuroimaging studies have identified distinct connectivity abnormalities across ACC subdivisions in MDD [2, 3]. More recently, increased connectivity between the sgACC and posterior cingulate cortex, alongside greater coupling with prefrontal and limbic regions, has been observed in antidepressant responders relative to non-responders [24]. Together, these findings suggest that anatomically specific ACC circuits may contribute differently to antidepressant response and warrant investigation of their changes during treatment.

Neuroimaging studies have demonstrated that antidepressant treatment is associated with changes in corticolimbic connectivity and functional organization of large-scale networks, including the default mode, salience, and executive-control networks [10, 11, 25]. However, these changes are heterogeneous, with some associated with symptom improvement or remission, whereas others occur regardless of clinical outcome or differ across treatment modalities [11, 12, 26]. These findings suggest that treatment-related changes in functional connectivity may not necessarily reflect neural processes associated with clinical improvement.

Complementary evidence from neuromodulation studies further highlights the clinical relevance of ACC-centered circuits. Specifically, antidepressant outcomes following prefrontal stimulation have been linked to intrinsic connectivity with the sgACC, while deep brain stimulation targeting the sgACC has implicated engagement of distributed pathways in sustained clinical improvement [27-29]. Together, these findings support a circuit-level understanding of antidepressant response but leave unresolved whether connectivity changes across distinct ACC subdivisions are specifically associated with clinical improvement.

In addition to the ACC, the hippocampal formation is a central region in depression. A meta-analysis identified increased resting-state connectivity between the default mode network and hippocampus in MDD [2], while a review highlighted broader abnormalities in connectivity within and between large-scale functional networks [30]. The hippocampus is critical to stress regulation [31], and has been implicated in MDD pathophysiology, including altered resting-state functional connectivity (rsFC) across hippocampal subfields [32]. Following 8 weeks of escitalopram treatment, increased bilateral hippocampal connectivity strength has also been reported [10], suggesting hippocampal connectivity changes could be another correlate of depression response.

In light of this prior evidence, the present study examined rsFC across 8 weeks of antidepressant treatment in individuals with MDD. Using anatomically defined seeds in the sgACC, pgACC, dACC, and hippocampal formation, we first tested overall pre- to-post rsFC changes and then examined whether changes differed by categorical response, remission, or dimensional improvement in depressive symptoms. We hypothesized that antidepressant treatment and clinical response would be associated with ACC subregion-specific changes in rsFC. Given heterogeneity in prior findings, we did not specify connection-level directional hypotheses.

## METHODS

### Participants and Procedures

The SMART Trial to Predict Anhedonia Response to Antidepressant Treatment (SMART-D) was a double-blinded, randomized controlled trial conducted at McLean Hospital in Belmont, MA, USA. All participants provided written informed consent to participate. SMART-D was designed to test a predesigned algorithm aimed to increase antidepressant treatment response to either sertraline or bupropion using pre-specified clinical, neural, and cognitive markers [33]. The current analyses included 41 treatment-seeking, unmedicated individuals with MDD who completed the study (i.e., provided clinical data up to week 8) and had usable pre- and post-treatment resting-state fMRI and clinical data. Participants were ages 18-65 (mean age 29.88 ± 9.37), met criteria for a current diagnosis of MDD assessed by the Structured Clinical Interview for DSM-5 Disorders, Clinician Version (SCID-5-CV) [34]. Participants were also required to have elevated depression symptoms, defined as a score >15 on the 17-item Hamilton Rating Scale for Depression (HAM-D-17) [35], and elevated anhedonia, defined as a score >24 on the 14-item Snaith–Hamilton Pleasure Scale (SHAPS) [36]. Participants with a lifetime diagnosis of bipolar disorder, psychotic disorders, current eating disorder, obsessive-compulsive disorder, post-traumatic stress disorder, substance use disorder, or resistant to more than 3 depression treatments (assessed by the Antidepressant Treatment Response Questionnaire) [37] were excluded from the screening session. In addition, participants with history of non-response to an adequate treatment with sertraline and bupropion in the current episode were excluded. Participants also needed to pass electrocardiogram screening to assess potential contraindication to bupropion or sertraline by study physicians. After screening, participants underwent baseline fMRI imaging session, assigned to biomarkers using the preregistered algorithm (for a detailed description of the bupropion- and sertraline-specific markers used to stratify treatment, see [33]), and were randomized to receive either bupropion or sertraline for 8 weeks. During the 8-week treatment, concomitant treatment with other psychotropic medications or change in psychotherapy were not allowed. After 8 weeks of antidepressant treatment, participants repeated the imaging session as post-treatment outcome assessment. 10-item Montgomery-Åsberg Depression Rating Scale (MADRS) [38] was administered by study clinician seven times (week 0, week 1-week 4, week 6, and week 8/post-treatment assessment) to monitor participant symptom improvement. Medication side-effects were self-reported using the Frequency, Intensity, and Burden of Side Effects Rating (FIBSER) [39] Scale and the Treatment-Emergent Signs and Symptoms (TESS) [40] checklist and assessed by study physician each week (see **Supplementary Table 1**).

Response to treatment was defined as a ≥50% reduction in MADRS scores from the pre-treatment session to the week 8, whereas participants with <50% reduction in MADRS score were classified as non-responders. Remission was defined as an outcome assessment MADRS score ≤10 [41], with scores >10 classified as non-remission.

### fMRI Data Acquisition and Processing

Structural and functional images were acquired on a 3-T Siemens Prisma-Fit system using a 32-channel head coil. A T1-weighted anatomical images were acquired using a three-dimensional MPRAGE sequence (repetition time of 2,400 ms, echo time of 2.22 ms, inversion time of 1,000 ms and flip angle of 8°) with 0.8-mm isotropic voxels (field of view of 256 mm, sagittal orientation, 208 slices). Parallel imaging used GRAPPA with a phase-encoding acceleration factor of 2; phase-encoding was anterior to posterior. A total of 300 BOLD images were acquired using a multi-echo gradient-echo echo planar imaging sequence (repetition time of 1.33 s, four echoes with echo times of 12.6, 29.16, 45.72 and 62.28 ms, flip angle of 67°, 60 axial slices, 2.5-mm isotropic voxels, multiband factor of 4, GRAPPA of 2, phase-encoding anterior to posterior and field of view of 216 mm × 216 mm). A brief reverse-phase (posterior to anterior) run in addition to the anterior-to-posterior run was acquired for TOPUP-based distortion correction.

The CONN toolbox pipeline (version 21.a, MATLAB R2022a) was used for processing. First, we ran the meica.py on the four individual echo trains to obtain the optimally combined echo images. We then ran the default CONN pipeline. Specifically, CONN uses SPM12 and includes realignment, normalization in Montreal Neurological Institute (MNI) space and smoothing with a 6-mm kernel. For motion correction and denoising, CONN uses SPM12 to assess head motion by translation and rotation in x, y and z directions. It then leverages artifact detection tools to calculate time points of significant head motion with more than 0.5 mm motion compared with the previous frame, or global mean intensity change of more than 3 S.D. from mean intensity across functional scans for each participant. Any participant with >15% outlier volumes out of the resting-state scan series was excluded from group-level analyses. Default CONN parameters were used for the remainder of the pipeline.

Resting-state functional connectivity (rsFC) analyses examined pre- to post-treatment changes in connectivity using *a priori* ACC and hippocampal seed regions defined from the Human Connectome Project (HCP) parcellation. ACC seeds included the sgACC (Brodmann area [BA] 25), pgACC (Glasser parcels a24 and p32), and dACC (parcels p24 and a32pr). The hippocampal formation seed comprised four anatomically distinct regions: the presubiculum (PreS), entorhinal cortex (EC), perirhinal/ectorhinal cortex (PeEC), and hippocampus. Functional connectivity was calculated between each of these regions and cortical regions across the brain. For each cortical region, connectivity values were then averaged across the four hippocampal formation regions to derive a composite hippocampal formation connectivity measure.

### Statistical analysis

Linear mixed-effects models were conducted to characterize overall treatment-related changes separately for each seed with cortical regions. rsFC was modeled as a function of timepoint (baseline, post-treatment), adjusting for age, sex, and mean framewise displacement (FD), with a subject-specific random intercept to account for repeated observations.

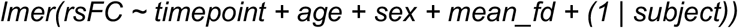

Associations between changes in rsFC from pre- to post-treatment and treatment outcomes were examined using both categorical and dimensional measures of clinical improvement. For categorical outcomes, separated mixed-effects models tested timepoint × treatment response and timepoint × remission status interactions, adjusting for age, sex, and mean FD and including a subject-specific random intercept.

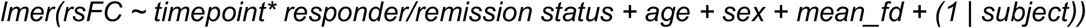

For dimensional analyses, associations between changes in rsFC and the magnitude of depressive symptom improvement were examined using MADRS percent change as a continuous measure.

Models tested the timepoint × MADRS percent-change interaction, adjusting for age, sex, and mean FD and including a subject-specific random intercept.

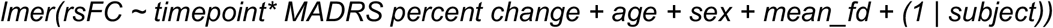

For each of these analyses, multiple-comparison correction was performed separately for each of the 4 seed regions (sgACC, pgACC, dACC, hippocampal formation) and hemisphere. Within each seed and hemisphere, false discovery rate (FDR) correction was applied across the 180 cortical regions. Results surviving FDR correction (*P*_FDR_ < .05) were considered statistically significant.

### Sensitivity analyses

Medication received and marker assignment (i.e., BUP, SERT, Both Markers, Neither) were separately included as covariates in sensitivity analyses to determine whether observed response-related effects were independent of treatment and marker assignment. Post hoc contrasts were conducted using estimated marginal means to characterize pre- to-post changes in rsFC separately for responders vs. non-responders and remitters vs. non-remitters. Differences in connectivity changes between groups were evaluated using difference-in-differences contrasts. Post hoc contrasts were evaluated using unadjusted p-values (p <.05). Significant timepoint × clinical outcome interactions were followed up with two-sided Fisher’s exact tests to compare the proportions of participants showing increased versus decreased rsFC between responders and non-responders or remitters and non-remitters. Odds ratios were calculated as effect-size estimates. Additionally, two-sided exact binomial tests were conducted to determine whether the proportion of participants exhibiting increased versus decreased connectivity within each clinical outcome group differed from an expected proportion of 50%.

## RESULTS

### Participants

Forty-one participants completed all study sessions (i.e., completed the 8-week clinical assessments) and had available imaging data (Table 1). Of these, 30 (73.2%) were classified as responders and 11 (26.8%) as non-responders, while 26 (63.4%) met criteria for remission and 15 (36.6%) were non-remitters. Participants had a mean age of 29.88 years (SD = 9.37), and 28 (68.3%) were female. The mean baseline MADRS score was 27.78 (SD = 6.35), with 6 participants (14.6%) classified as having mild, 30 (73.2%) moderate, and 5 (12.2%) severe depressive symptoms. Demographic and clinical characteristics stratified by treatment response and remission status are presented in Table 1.

**TABLE 1.** Baseline demographics and clinical characteristics for SMART-D participants who completed 8 weeks of treatment as well as both pre- and post-treatment MRI scans.

|  |  | **Treatment Response** |  |
|---|---|---|---|
|  | **Overall<br>(N=41)** | **Non-Responders<br>(N=11)** | **Responders<br>(N=30)** |
| Age, years | 29.88 (9.37) | 26.82 (10.76) | 31.00 (8.73) |
| Female | 28 (68.3%) | 8 (72.7%) | 20 (66.7%) |
| Hispanic/Latino | 5 (13.2%) | 2 (20.0%) | 3 (10.7) |
| Race |  |  |  |
| White | 20 (54.1%) | 4 (44.4%) | 16 (57.1%) |
| Black | 4 (10.8%) | 0 (0.0%) | 4 (14.3%) |
| Asian | 9 (24.3%) | 4 (44.4%) | 5 (17.9%) |
| Other race | 2 (5.4%) | 0 (0.0%) | 2 (7.1%) |
| Native American | 0 (0.0%) | 0 (0.0%) | 0 (0.0%) |
| More than one race | 2 (4.9%) | 1 (9.1%) | 1 (3.3%) |
| Education, years | 15.81 (2.83) | 15.50 (3.42) | 15.92 (2.69) |
| Employed | 21 (70.0%) | 5 (71.4%) | 16 (69.6%) |
| Baseline MADRS | 27.78 (6.35) | 28.64 (6.64) | 27.47 (6.33) |
| Baseline SHAPS | 36.54 (5.84) | 37.18 (7.95) | 36.30 (5.00) |
| Age of MDD first onset, years | 20.17 (8.25) | 20.45 (8.79) | 20.07 (8.20) |
| Number of lifetime depressive episodes | 7.05 (11.61) | 3.73 (4.03) | 8.31 (13.26) |

*Note.* Continuous variables are presented as *M* (*SD*); categorical variables are presented as *n* (%). MADRS = Montgomery-Åsberg Depression Rating Scale. Response to treatment: Responders: >50% improvement in MADRS from baseline to week 8. Remission status: Remitters: week 8 MADRS score ≤ 10

### Changes in Functional Connectivity Following Treatment

Across all participants (i.e., irrespective of clinical outcome), treatment was associated with significant changes in dACC and pgACC rsFC. Specifically, dACC connectivity decreased with regions of the default mode and frontoparietal networks from pre- to post-treatment (**Figure 1A**, 9 regions significant after FDR correction, 0.002 < *P*_FDR_< 0.041). We also observed pgACC connectivity decreased across multiple cortical regions (**Figure 1B**, 15 regions significant after FDR correction, 0.005 < *P*_FDR_< 0.049). No significant overall change was observed for sgACC or hippocampal formation connectivity (*P*_FDR_ > .05) (**Figure 1C, D**).

**Figure 1.**
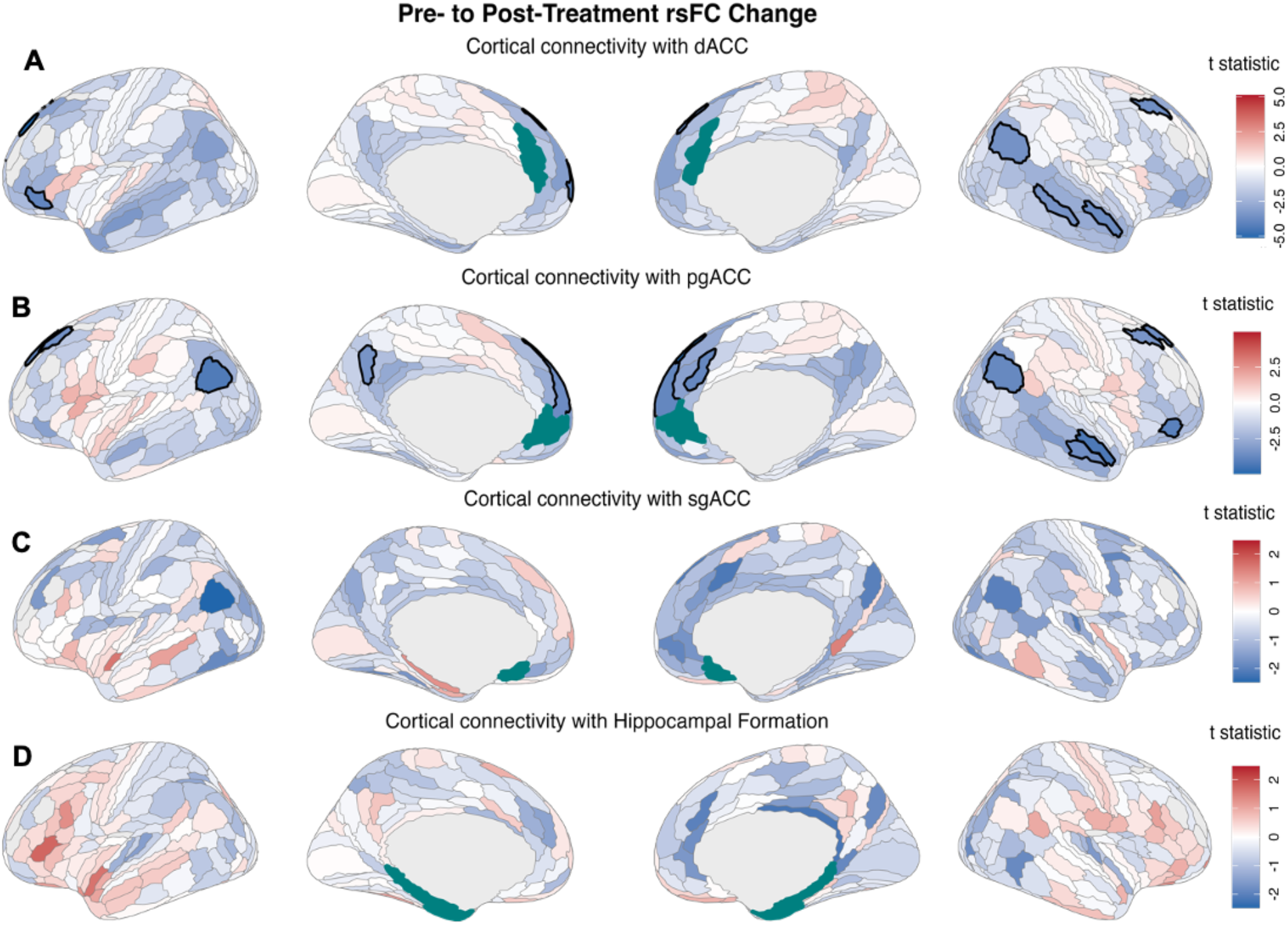
Changes in rsFC following 8 weeks of antidepressant treatment. Whole-brain cortical maps show *t*-statistics for pre- to post-treatment changes in rsFC with **(A)** dorsal anterior cingulate cortex (dACC, parcels p24&a32pr), **(B)** pregenual anterior cingulate cortex (pgACC, parcels a24&p32), **(C)** subgenual anterior cingulate cortex (sgACC, BA25), and **(D)** hippocampal formation. Warm colors indicate positive *t*-statistics and cool colors indicate negative *t*-statistics. Black outlines denote connections surviving false discovery rate correction (*P*_FDR_ < .05). Seed regions are shown in teal.

### Functional Connectivity Changes Associated With Treatment Response

Changes in sgACC connectivity differed significantly between responders and non-responders (**Figure 2A, B**; 9 regions significant after FDR correction, 0.005 < *P*_FDR_ < 0.049). These regions were primarily distributed across medial and lateral temporal and inferior frontal regions. The strongest response-related differences were observed for connectivity with the right posterior middle/inferior temporal cortex (TE1p), t(33.78) = 4.87, *P*_FDR_ = .005, and right posterior inferior frontal sulcus (IFSp), t(35.59) = 4.33, *P*_FDR_ = .011. Significant differences were also observed for right posterior portion of area 47r in the inferior frontal gyrus (p47r), t(32.70) = 3.40, *P*_FDR_ = .045. Post hoc estimated marginal mean contrasts revealed opposing patterns of connectivity change between groups, with non-responders generally exhibiting decreases and responders showing increases in connectivity. For example, right TE1p showed decreased connectivity among non-responders (estimated change = −1.50) and increased connectivity among responders (estimated change = 0.98). A similar pattern was observed for right p47r, t(32.70) = 3.40, *P*_FDR_ = .045, with significant decreases among non-responders (estimated change = −1.14, *p* = .016) and increases among responders (estimated change = 0.69, *p* = .025). Individual trajectories for these two representative connections further illustrate the divergent patterns of connectivity change between groups (**Figure 2C**). Consistent with these findings, Fisher’s exact tests revealed significant differences in the directions of connectivity change between groups for the right TE1p (*p* = .002) and right p47r (*p* = .003). For right TE1p, 24 of 30 responders (80.0%; binomial *p* = .007) exhibited increased connectivity, compared with 2 of 11 non-responders (18.2%; binomial *p* = .065). Similarly, for right p47r, 23 of 30 responders (76.0%; binomial *p* = .023) showed increased connectivity, compared with 2 of 11 non-responders (18.2%; binomial *p* = .065).

**Figure 2.**
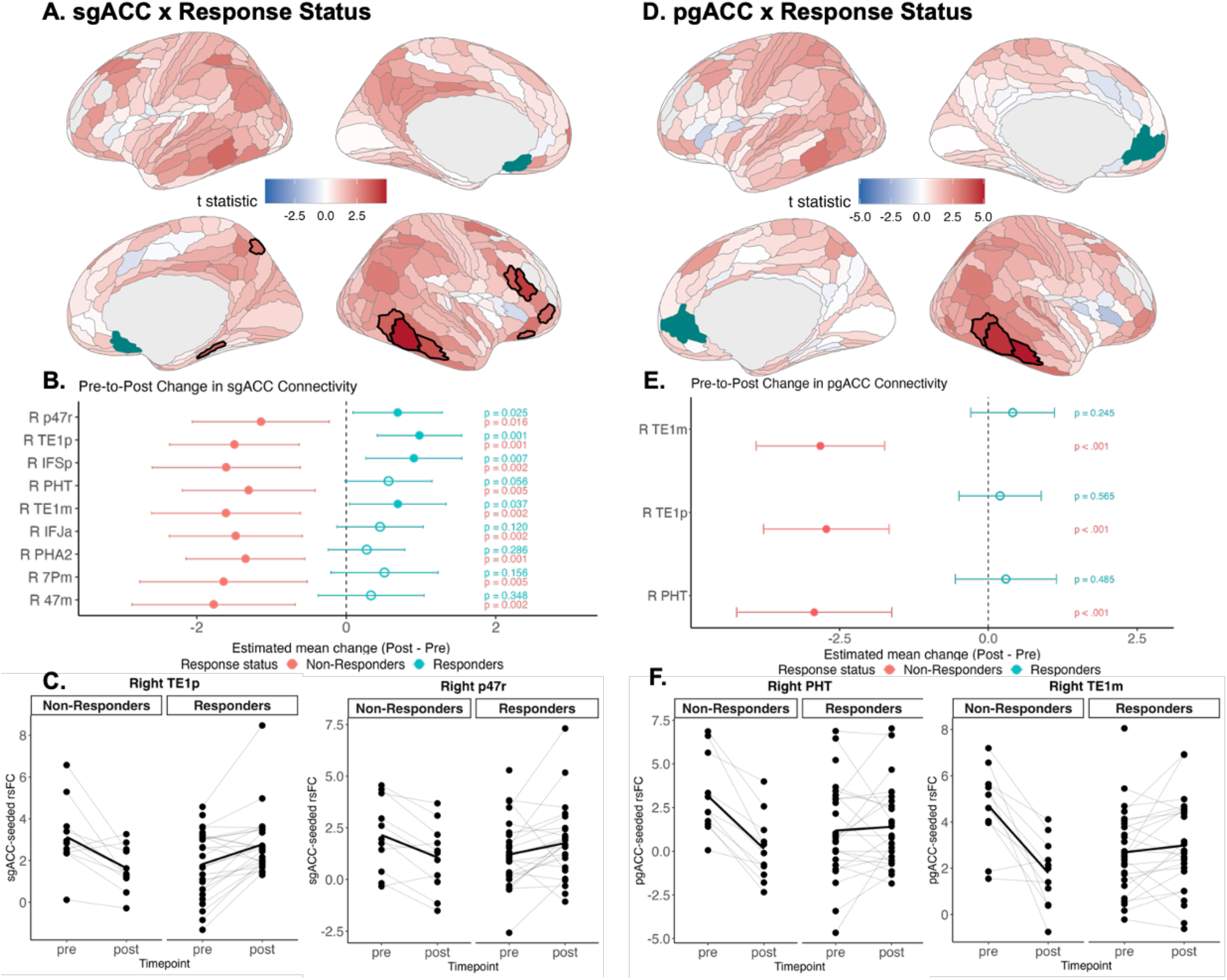
Changes in rsFC associated with response to antidepressant treatment. **(A)** Whole-brain cortical maps show t-statistics for differences in pre- to post-treatment resting-state functional connectivity (rsFC) changes between responders and non-responders for the subgenual anterior cingulate cortex (sgACC) seed. Warm colors indicate positive t-statistics, whereas cool colors indicate negative t-statistics. Black outlines denote regions surviving false discovery rate (FDR) correction (*P*_FDR_ < .05), and the sgACC seed is shown in teal. **(B)** Forest plots display model-estimated pre- to post-treatment rsFC changes and 95% confidence intervals for non-responders (coral) and responders (teal) across regions showing significant response-related differences in sgACC connectivity changes. Corresponding p-values indicate the significance of within-group pre- to post-treatment changes. **(C)** Spaghetti plots illustrate individual pre- to post-treatment rsFC trajectories for representative sgACC connections, stratified by treatment response status. Gray lines represent individual trajectories, and bold black lines represent group means. **(D)** Whole-brain cortical maps show t-statistics for response-related differences in pre- to post-treatment rsFC changes for the pregenual anterior cingulate cortex (pgACC) seed, using the same color conventions as in panel A. **(E)** Forest plots display model-estimated pre- to post-treatment rsFC changes and 95% confidence intervals for non-responders (coral) and responders (teal) across regions showing significant response-related differences in pgACC connectivity changes. Corresponding p-values indicate the significance of within-group pre- to post-treatment changes. **(F)** Spaghetti plots illustrate individual pre- to post-treatment rsFC trajectories for representative pgACC connections, stratified by treatment response status. Gray lines represent individual trajectories, and bold black lines represent group means.

Similarly, significant response-related differences in pgACC connectivity changes were observed for three right temporal regions: the middle portion of the middle temporal gyrus (TE1m), t(30.47) = 5.09, *P*_FDR_ = .003, posterior middle/inferior temporal cortex (TE1p), t(31.73) = 4.71, *P*_FDR_ = .004, and posterior temporal cortex (PHT), t(34.26) = 4.21, *P*_FDR_ = .010 (**Figure 2D, E**). Post hoc estimated marginal mean contrasts revealed decreased connectivity among non-responders and increased connectivity among responders for right TE1m (−2.82 vs. 0.41, *p* < .001), TE1p (−2.72 vs. 0.20, *p* < .001), and PHT (−2.93 vs. 0.30, *p* < .001). Individual trajectories for representative connections are shown in **Figure 2F**. Consistent with these findings, Fisher’s exact tests revealed significant differences in the direction of connectivity change between responders and non-responders for right TE1m (*p* < .001), TE1p (*p* = .009), and PHT (*p* = .005). For right TE1m, 21 of 30 responders (70.8%; binomial *p* = .064) exhibited increased connectivity, compared with 1 of 11 non-responders (9.1%; binominal *p* = .012). Similarly, increased connectivity was observed in 17 of 30 responders (58.3%; binominal *p* = .541) vs. 1 of 11 non-responders (9.1%; binominal *p* = .012) for right TE1p, and 12 of 24 responders (50.0%; binomial *p* = 1.000) vs. none of the 11 non-responders (0%; binomial *p* < .001) for right PHT. No response-related differences survived FDR correction for the dACC or hippocampal formation seeds (*P*_FDR_ > .05).

### Functional Connectivity Changes Associated With Remission

Changes in sgACC connectivity differed between remitters and non-remitters for left dorsolateral prefrontal cortex (area 8Ad) and middle temporal gyrus (area TE1m; ts = 4.39–4.46, *P*_FDR_ = .009; **Figure 3A**). Post hoc estimated marginal mean contrasts showed opposing patterns of connectivity change between groups, with decreased connectivity among non-remitters and increased connectivity among remitters for left 8Ad (−1.62 vs. 0.60, *p* < .001) and TE1m (−1.48 vs. 0.84, *p* < .001). Individual trajectories for these two connections further illustrate the divergent patterns of connectivity change between groups (**Figure 3B**). Fisher’s exact tests confirmed significant differences in the direction of change for sgACC connectivity with left 8Ad (*p* < .001) and left TE1m (*p* = .002). For left 8Ad, 17 of 26 remitters (65.0%; binomial *p* = .263) exhibited increased connectivity, compared with 1 of 15 non-remitters (6.7%; binomial *p* <.001). Similarly, increased connectivity with left TE1m was observed in 18 of 26 remitters (70%; binomial *p* = .115), compared with 2 of 15 non-remitters (13.3%; binomial *p* = .007). No significant remission-related changes were observed for pgACC, dACC, or hippocampal formation seeds (*P*_FDR_ > .05).

**Figure 3.**
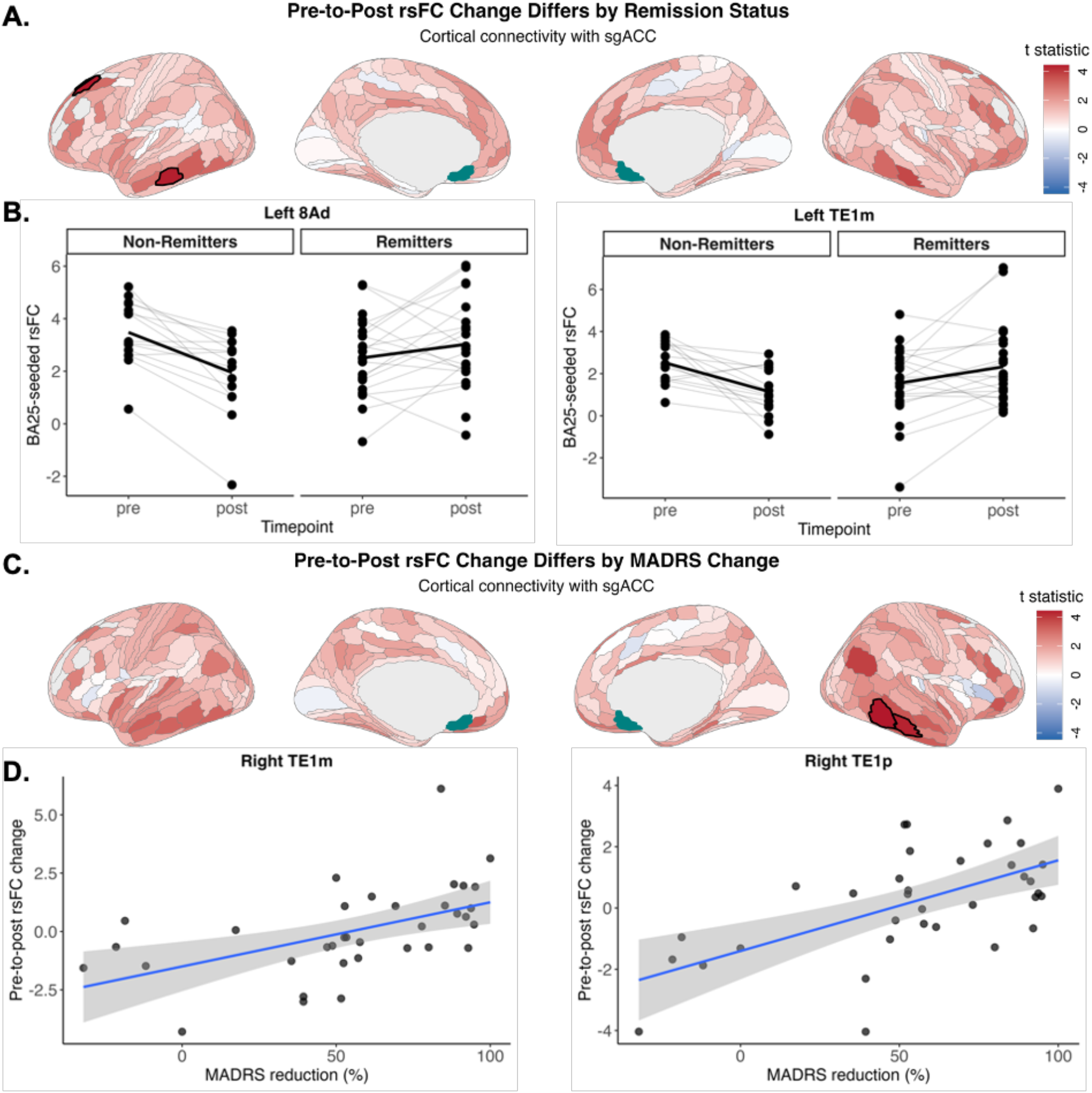
Changes in sgACC resting-state functional connectivity associated with antidepressant treatment outcomes. **(A)** Whole-brain cortical maps show t-statistics for associations between pre- to post-treatment changes in subgenual anterior cingulate cortex (sgACC) resting-state functional connectivity (rsFC) and remission status. Warm colors indicate positive t-statistics, whereas cool colors indicate negative t-statistics. Black outlines denote regions surviving false discovery rate (FDR) correction (*P*_FDR_ < .05), and the sgACC seed is shown in teal. **(B)** Plots illustrate individual pre- to post-treatment rsFC trajectories for significant connections with left 8Ad and left TE1m, stratified by remission status. Gray lines represent individual trajectories, and bold black lines represent group means. **(C)** Whole-brain cortical maps show t-statistics for associations between pre- to post-treatment changes in sgACC rsFC and percent reduction in Montgomery–Åsberg Depression Rating Scale (MADRS) scores. **(D)** Scatterplots illustrate associations between percent MADRS reduction and pre- to post-treatment rsFC changes for right TE1m and right TE1p. Blue lines represent fitted linear regressions, with shaded areas indicating 95% confidence intervals.

### Association Between Connectivity and Symptom Improvement

Corroborating categorical analyses, greater percent reductions in MADRS scores were significantly associated with pre- to post-treatment changes in sgACC connectivity with right TE1p and TE1m (*P*_FDR_ = .023, **Figure 3C, D**). These associations indicate that greater improvement in depressive symptoms was accompanied by greater increases in sgACC connectivity with temporal cortical regions. Individual associations for right TE1m and TE1p are illustrated in **Figure 3D**. No significant associations with MADRS improvement were observed for pgACC, dACC, or hippocampal formation seeds (*P*_FDR_ > .05).

### Sensitivity Analyses

Findings were unchanged after accounting for medication (bupropion vs. sertraline) received during the 8 weeks of treatment. Results were largely consistent after adjusting for marker assignment, with effect estimates remaining similar across analyses. However, several effects no longer survived FDR correction, particularly response-related sgACC and pgACC findings and overall pgACC dynamic treatment effects. Remission-related findings were largely unchanged. Full results are reported in **Supplementary Table 2**.

## DISCUSSION

This study examined changes in ACC and hippocampal formation rsFC after 8 weeks of monoaminergic antidepressant treatment, and their associations with clinical improvement in MDD. Irrespective of clinical outcome, antidepressant treatment was associated with widespread reductions in the dACC and pgACC connectivity, particularly with regions of the default mode and frontoparietal networks. The most prominent findings emerged when considering individual differences in treatment outcome. Specifically, treatment responders showed increases in sgACC and pgACC connectivity with temporal and inferior frontal regions, whereas non-responders predominantly showed decreased in these connections. Remission status was similarly associated with increased sgACC connectivity, and dimensional analyses also linked greater symptom improvement to sgACC connectivity change with temporal cortex. Thus, while dACC connectivity changed regardless of treatment outcomes, sgACC and pgACC connectivity tracked clinically relevant heterogeneity in antidepressant treatment response.

The most consistent outcome-related effects involved the sgACC. This region has long been implicated in MDD and antidepressant response through its interactions with distributed affective, reward, and regulatory systems [2, 23]. Previous studies have identified pretreatment ACC connectivity patterns associated with subsequent antidepressant outcomes, including differential responses to pharmacotherapy and psychotherapy [15, 17]. More recent evidence has also demonstrated the predictive relevance of pretreatment dACC connectivity across independent antidepressant trials [42]. Our findings extend this baseline predictor literature by examining treatment-related connectivity changes rather than baseline predictors alone. Although sgACC connectivity did not change significantly across the sample as a whole, responders and remitters were substantially more likely to show increases in specific sgACC connectivity, whereas poorer outcomes were characterized by decreases. Moreover, sgACC connectivity change was associated with symptom improvement when outcome was modeled dimensionally. The convergence across response, remission, and continuous symptom change suggests that sgACC connectivity captures information about clinical improvement that is not apparent from average treatment-related change. Interestingly, despite widespread treatment-related changes in dACC connectivity, no significant dACC changes were associated with clinical outcomes in the primary analyses. This distinction may also help explain heterogeneity across previous studies reporting different directions or magnitudes of sgACC change following depression treatment.

The anatomical distribution of these findings further highlights the relevance of interactions between ACC subdivisions and distributed temporal and frontal cortical regions. Response-related differences in sgACC and pgACC connectivity involved temporal regions (e.g. TE1m and TE1p), with additional sgACC effects involving inferior frontal cortex and pgACC effects involving posterior temporal cortex. Remission-related sgACC differences similarly involved temporal and dorsal prefrontal regions. These findings are consistent with previous evidence implicating sgACC-temporal connectivity in depression treatment outcomes. For example, greater pre-treatment connectivity between the sgACC and middle temporal gyrus was associated with subsequent symptom improvement during psychotherapy for late-life depression [43]. Although the present study examined longitudinal changes rather than baseline connectivity, both findings suggest that the clinical relevance of sgACC circuitry extends beyond its interactions with traditionally emphasized limbic and prefrontal regions. The involvement of temporal cortical connections across several outcome measures may reflect broader changes in the functional integration of sgACC-centered networks during clinical improvement. However, the specific psychological processes underlying these associations remain unclear, and the observed connectivity changes cannot establish the direction of functional influence between regions.

The dissociation between overall treatment-related changes and those associated with clinical improvement represents another important finding. Across participants, the predominant pattern was decreased dACC and pgACC connectivity, whereas favorable treatment outcomes were associated with relatively greater increases or smaller decreases in selected sgACC and pgACC connections. This distinction is consistent with emerging evidence that antidepressant-associated connectivity changes are heterogeneous and may reflect multiple processes rather than a uniform neural signature of recovery. For example, previous work identified both shared and treatment-specific connectivity changes among patients achieving remission following pharmacotherapy and cognitive behavioral therapy, with medication-related remission characterized predominantly by reductions in connectivity across several networks [11]. Similarly, early changes in dynamic resting-state network organization have been associated with sertraline response and subsequent symptom improvement [44]. More recently, distinct patterns of early connectivity change have been identified across antidepressant- and placebo-treated patients, including changes occurring independently of treatment outcome and others associated with placebo or medication-specific effects [12]. Together, these studies highlight the importance of distinguishing neural changes occurring during treatment from those specifically associated with clinical outcomes. By explicitly examining timepoint-by-outcome interactions across anatomically distinct ACC subdivisions, the present study identified connectivity trajectories that were not captured by overall pre- to post-treatment analyses. These findings suggest that the neural correlates of successful antidepressant response may be more anatomically specific than the broader connectivity changes observed across treated patients.

Several limitations should be considered. First, although unmedicated and all participants completed the 8-week treatment, the sample was relatively small (*N*=41), particularly the non-responder subgroup, limiting power to characterize heterogeneity in connectivity change. Nevertheless, convergent effects were observed across categorical response, remission, and dimensional MADRS improvement, providing support for an association between the ACC connectivity change and clinical outcome. Several response-related sgACC and pgACC effects no longer survived FDR correction after adjustment for marker assignment, although effect estimates remained similar, underscoring the need for replication in larger samples and for adequately powered analyses of medication-specific effects. Second, rsFC was assessed only before and after treatment. Thus, the present study cannot determine when these connectivity changes emerge during treatment or whether they precede, accompany, or follow clinical improvement. More frequent imaging with clinical assessment during treatment could establish the temporal sequence of these changes and determine whether ACC connectivity shift precedes subsequent clinical response.

In sum, a convergence of categorical and dimensional analyses implicates sgACC connectivity change as a feature of successful antidepressant response rather than a general consequence of treatment exposure. More generally, these findings suggest that improvement during antidepressant treatment involves a pattern of ACC circuit reorganization that is distinguishable from the broader connectivity changes occurring across treated patients. Establishing when these outcome-related changes emerge, and whether they prospectively identify patients likely to benefit from treatment, will be important for determining their utility as dynamic markers of antidepressant response.

## Acknowledgements

Work on ‘McLean Dissection of Anhedonia: From Neural Systems and Multiomics to Behavior’ was supported by Wellcome Leap as part of the Multi-Channel Psych Program (awarded to D.A.P.). D.A.P. was partially supported by grant R37 MH068376. The funders had no role in the design and conduct of the study; collection, management, analysis and interpretation of the data; preparation, review or approval of the manuscript; or the decision to submit the manuscript for publication. The content is solely the responsibility of the authors and does not necessarily represent the official views of the National Institutes of Health or Wellcome Leap. We thank the participants of the SMART-D study for their time and contributions to this research.

## References

1. Hasin, D. S., Sarvet, A. L., Meyers, J. L., Saha, T. D., Ruan, W. J., Stohl, M., & Grant, B. F. (2018). Epidemiology of adult DSM-5 major depressive disorder and its specifiers in the United States. JAMA Psychiatry, 75(4), 336–346. 10.1001/jamapsychiatry.2017.4602

2. Kaiser, R. H., Andrews-Hanna, J. R., Wager, T. D., & Pizzagalli, D. A. (2015). Large-scale network dysfunction in major depressive disorder: A meta-analysis of resting-state functional connectivity. JAMA Psychiatry, 72(6), 603–611. 10.1001/jamapsychiatry.2015.0071

3. Zhou, Z., Gao, Y., Bao, W., Liang, K., Cao, L., Tang, M., Li, H., Hu, X., Zhang, L., Sun, H., Roberts, N., Gong, Q., & Huang, X. (2024). Distinctive intrinsic functional connectivity alterations of anterior cingulate cortex subdivisions in major depressive disorder: A systematic review and meta-analysis. Neuroscience & Biobehavioral Reviews, 159, Article 105583. 10.1016/j.neubiorev.2024.105583

4. Drevets, W. C., Savitz, J., & Trimble, M. (2008). The subgenual anterior cingulate cortex in mood disorders. CNS Spectrums, 13(8), 663–681. 10.1017/S1092852900013754

5. Dunlop, B. W., Cha, J., Choi, K. S., Nemeroff, C. B., Craighead, W. E., & Mayberg, H. S. (2023). Functional connectivity of salience and affective networks among remitted depressed patients predicts episode recurrence. Neuropsychopharmacology, 48(13), 1901–1909. 10.1038/s41386-023-01653-w

6. MacQueen, G. M., Yucel, K., Taylor, V. H., Macdonald, K., & Joffe, R. T. (2008). Posterior hippocampal volumes are associated with remission rates in patients with major depressive disorder. Biological Psychiatry, 64(10), 880–883. 10.1016/j.biopsych.2008.06.027

7. Whitton, A. E., Treadway, M. T., & Pizzagalli, D. A. (2015). Reward processing dysfunction in major depression, bipolar disorder and schizophrenia. Current Opinion in Psychiatry, 28(1), 7–12. 10.1097/YCO.0000000000000122

8. Dichter, G. S., Gibbs, D., & Smoski, M. J. (2015). A systematic review of relations between resting-state functional-MRI and treatment response in major depressive disorder. Journal of Affective Disorders, 172, 8–17. 10.1016/j.jad.2014.09.028

9. Chai, Y., Sheline, Y. I., Oathes, D. J., Balderston, N. L., Rao, H., & Yu, M. (2023). Functional connectomics in depression: Insights into therapies. Trends in Cognitive Sciences, 27(9), 814–832. 10.1016/j.tics.2023.05.006

10. Wang, L., Xia, M., Li, K., Zeng, Y., Su, Y., Dai, W., et al. (2015). The effects of antidepressant treatment on resting-state functional brain networks in patients with major depressive disorder. Human Brain Mapping, 36(2), 768–778. 10.1002/hbm.22663

11. Dunlop, B. W., Cha, J., Choi, K. S., Rajendra, J. K., Nemeroff, C. B., Craighead, W. E., & Mayberg, H. S. (2023). Shared and unique changes in brain connectivity among depressed patients after remission with pharmacotherapy versus psychotherapy. American Journal of Psychiatry, 180(3), 218–229. 10.1176/appi.ajp.21070727

12. Tong, X., Fonzo, G. A., Carlisle, N. B., Xie, H., Berdichevsky, Y., Keller, C. J., et al. (2026). Early brain functional connectivity changes induced by antidepressants and placebo. Nature Mental Health. 10.1038/s44220-026-00729-y

13. Rush, A. J., Trivedi, M. H., Wisniewski, S. R., Nierenberg, A. A., Stewart, J. W., Warden, D., et al. (2006). Acute and longer-term outcomes in depressed outpatients requiring one or several treatment steps: A STAR*D report. American Journal of Psychiatry, 163(11), 1905–1917. 10.1176/ajp.2006.163.11.1905

14. Trivedi, M. H., Rush, A. J., Wisniewski, S. R., Nierenberg, A. A., Warden, D., Ritz, L., et al. (2006). Evaluation of outcomes with citalopram for depression using measurement-based care in STAR*D: Implications for clinical practice. American Journal of Psychiatry, 163(1), 28–40. 10.1176/appi.ajp.163.1.28

15. Dunlop, B. W., Rajendra, J. K., Craighead, W. E., Kelley, M. E., McGrath, C. L., Choi, K. S., et al. (2017). Functional connectivity of the subcallosal cingulate cortex and differential outcomes to treatment with cognitive-behavioral therapy or antidepressant medication for major depressive disorder. American Journal of Psychiatry, 174(6), 533–545. 10.1176/appi.ajp.2016.16050518

16. Whitton, A. E., Webb, C. A., Dillon, D. G., Kayser, J., Rutherford, A., Goer, F., et al. (2019). Pretreatment rostral anterior cingulate cortex connectivity with salience network predicts depression recovery: Findings from the EMBARC randomized clinical trial. Biological Psychiatry, 85(10), 872–880. 10.1016/j.biopsych.2018.12.007

17. Ang, Y.-S., Kaiser, R., Deckersbach, T., Almeida, J., Phillips, M. L., Chase, H. W., et al. (2020). Pretreatment reward sensitivity and frontostriatal resting-state functional connectivity are associated with response to bupropion after sertraline nonresponse. Biological Psychiatry, 88(8), 657–667. 10.1016/j.biopsych.2020.04.009

18. Korgaonkar, M. S., Goldstein-Piekarski, A. N., Fornito, A., & Williams, L. M. (2020). Intrinsic connectomes are a predictive biomarker of remission in major depressive disorder. Molecular Psychiatry, 25(7), 1537–1549. 10.1038/s41380-019-0574-2

19. Wang, Y., Wang, C., Zhou, J., Chen, X., Liu, R., Zhang, Z., et al. (2024). Contribution of resting-state functional connectivity of the subgenual anterior cingulate to prediction of antidepressant efficacy in patients with major depressive disorder. Translational Psychiatry, 14, Article 399. 10.1038/s41398-024-03117-1

20. Vogt, B. A. (2005). Pain and emotion interactions in subregions of the cingulate gyrus. Nature Reviews Neuroscience, 6(7), 533–544. 10.1038/nrn1704

21. Margulies, D. S., Kelly, A. M. C., Uddin, L. Q., Biswal, B. B., Castellanos, F. X., & Milham, M. P. (2007). Mapping the functional connectivity of anterior cingulate cortex. NeuroImage, 37(2), 579–588. 10.1016/j.neuroimage.2007.05.019

22. Beckmann, M., Johansen-Berg, H., & Rushworth, M. F. S. (2009). Connectivity-based parcellation of human cingulate cortex and its relation to functional specialization. Journal of Neuroscience, 29(4), 1175–1190. 10.1523/JNEUROSCI.3328-08.2009

23. Pizzagalli, D. A. (2011). Frontocingulate dysfunction in depression: Toward biomarkers of treatment response. Neuropsychopharmacology, 36(1), 183–206. 10.1038/npp.2010.166

24. Kang, Y., Jung, J., Han, K.-M., Auer, D. P., & Ham, B.-J. (2026). Resting-state functional connectivity of the fronto-limbic and default mode network as neural correlates of antidepressant response in major depressive disorder. Journal of Affective Disorders, 394, Article 120607. 10.1016/j.jad.2025.120607

25. Anand, A., Li, Y., Wang, Y., Wu, J., Gao, S., Bukhari, L., et al. (2005). Antidepressant effect on connectivity of the mood-regulating circuit: An fMRI study. Neuropsychopharmacology, 30(7), 1334–1344. 10.1038/sj.npp.1300725

26. Ju, Y., Wang, M., Liu, J., Liu, B., Yan, D., Lu, X., et al. (2023). Modulation of resting-state functional connectivity in default mode network is associated with the long-term treatment outcome in major depressive disorder. Psychological Medicine, 53(13), 5963–5975. 10.1017/S0033291722002628

27. Fox, M. D., Buckner, R. L., White, M. P., Greicius, M. D., & Pascual-Leone, A. (2012). Efficacy of transcranial magnetic stimulation targets for depression is related to intrinsic functional connectivity with the subgenual cingulate. Biological Psychiatry, 72(7), 595–603. 10.1016/j.biopsych.2012.04.028

28. Riva-Posse, P., Choi, K. S., Holtzheimer, P. E., Crowell, A. L., Garlow, S. J., Rajendra, J. K., et al. (2018). A connectomic approach for subcallosal cingulate deep brain stimulation surgery: Prospective targeting in treatment-resistant depression. Molecular Psychiatry, 23(4), 843–849. 10.1038/mp.2017.59

29. Siddiqi, S. H., Kandala, S., Hacker, C. D., Trapp, N. T., Leuthardt, E. C., Carter, A. R., & Brody, D. L. (2023). Individualized precision targeting of dorsal attention and default mode networks with rTMS in traumatic brain injury-associated depression. Scientific Reports, 13, Article 4052. 10.1038/s41598-022-21905-x

30. Mulders, P. C., van Eijndhoven, P. F., Schene, A. H., Beckmann, C. F., & Tendolkar, I. (2015). Resting-state functional connectivity in major depressive disorder: A review. Neuroscience & Biobehavioral Reviews, 56, 330–344. 10.1016/j.neubiorev.2015.07.014

31. McEwen, B. S., Nasca, C., & Gray, J. D. (2016). Stress effects on neuronal structure: Hippocampus, amygdala, and prefrontal cortex. Neuropsychopharmacology, 41, 3–23. 10.1038/npp.2015.171

32. Hao, Z. Y., Zhong, Y., Ma, Z. J., Xu, H. Z., Kong, J. Y., Wu, Z., et al. (2020). Abnormal resting-state functional connectivity of hippocampal subfields in patients with major depressive disorder. BMC Psychiatry, 20, Article 71. 10.1186/s12888-020-02490-7

33. Zhukovsky, P., Kuhn, M., Borchers, L. R., et al. (2026). A precision medicine trial of bupropion and sertraline for major depressive disorder using a biomarker-guided sequential multiple-assignment design. Nature Mental Health, 4, 1099–1108. 10.1038/s44220-026-00671-z

34. First, M. B., Williams, J. B. W., Karg, R. S., & Spitzer, R. L. (2016). Structured Clinical Interview for DSM-5 Disorders, Clinician Version (SCID-5-CV). American Psychiatric Association Publishing.

35. Hamilton, M. (1960). A rating scale for depression. Journal of Neurology, Neurosurgery & Psychiatry, 23(1), 56–62. 10.1136/jnnp.23.1.56

36. Snaith, R. P., Hamilton, M., Morley, S., Humayan, A., Hargreaves, D., & Trigwell, P. (1995). A scale for the assessment of hedonic tone the Snaith-Hamilton Pleasure Scale. British Journal of Psychiatry, 167(1), 99–103. 10.1192/bjp.167.1.99

37. Chandler, G. M., Iosifescu, D. V., Pollack, M. H., Targum, S. D., & Fava, M. (2010). Validation of the Massachusetts General Hospital Antidepressant Treatment History Questionnaire (ATRQ). CNS Neuroscience & Therapeutics, 16(5), 322–325. 10.1111/j.1755-5949.2009.00102.x

38. Montgomery, S. A., & Åsberg, M. (1979). A new depression scale designed to be sensitive to change. British Journal of Psychiatry, 134(4), 382–389. 10.1192/bjp.134.4.382

39. Wisniewski, S. R., Rush, A. J., Balasubramani, G. K., Trivedi, M. H., & Nierenberg, A. A. (2006). Self-rated global measure of the frequency, intensity, and burden of side effects. Journal of Psychiatric Practice, 12(2), 71–79. 10.1097/00131746-200603000-00002

40. National Institute of Mental Health (NIMH). (1976). Treatment Emergent Symptom Scale (TESS). In W. Guy (Ed.), ECDEU assessment manual for psychopharmacology (pp. 322–347). Rockville, MD: U.S. Department of Health, Education, and Welfare. (DHEW Publication No. ADM 76-338).

41. Zimmerman, M., Posternak, M. A., & Chelminski, I. (2004). Derivation of a definition of remission on the Montgomery–Asberg depression rating scale corresponding to the definition of remission on the Hamilton rating scale for depression. Journal of Psychiatric Research, 38(6), 577–582. 10.1016/j.jpsychires.2004.03.007

42. Zhukovsky, P., Trivedi, M. H., Weissman, M., Parsey, R., Kennedy, S., & Pizzagalli, D. A. (2025). Generalizability of treatment outcome prediction across antidepressant treatment trials in depression. JAMA Network Open, 8(3), Article e251310. 10.1001/jamanetworkopen.2025.1310

43. Solomonov, N., Victoria, L. W., Dunlop, K., Respino, M., Hoptman, M. J., Zilcha-Mano, S., et al. (2020). Resting state functional connectivity and outcomes of psychotherapies for late-life depression. American Journal of Geriatric Psychiatry, 28(8), 859–868. 10.1016/j.jagp.2020.04.008

44. Kaiser, R. H., Chase, H. W., Phillips, M. L., Deckersbach, T., Parsey, R. V., Fava, M., et al. (2022). Dynamic resting-state network biomarkers of antidepressant treatment response. Biological Psychiatry, 92(7), 533–542. 10.1016/j.biopsych.2022.03.020

