## Supplemental Tables for "Changes in anterior cingulate connectivity associated with response to antidepressant treatment for major depressive disorder"

**Supplementary Information**

**Supplementary Table 1:** Side effects reported during 8 weeks of antidepressant treatment

|  | **No. (%)** | |
| --- | --- | --- |
| **Side Effect** | **Sertraline (N=21)** | **Bupropion (N=20)** |
| Insomnia | 14 (66.7%) | 10 (50.0) |
| Depressive affect | 11 (52.4%) | 11 (55.0) |
| Drowsiness | 13 (61.9%) | 7 (35.0) |
| Headache | 8 (38.1%) | 9 (45.0) |
| Dry mouth | 8 (38.1%) | 9 (45.0) |
| Excitement/agitation | 5 (23.8%) | 7 (35.0) |
| Increased motor activity | 5 (23.8%) | 6 (30.0) |
| Decreased appetite | 7 (33.3%) | 2 (10.0) |
| Nausea/vomiting | 7 (33.3%) | 4 (20.0) |
| Sweating | 5 (23.8%) | 3 (15.0) |
| Weight gain | 3 (14.3%) | 6 (30.0) |
| Decreased motor activity | 4 (19.0%) | 3 (15.0) |
| Constipation | 6 (28.6%) | 4 (20.0) |
| Diarrhea | 2 (9.5%) | 1 (5.0) |
| Nasal congestion | 2 (9.5%) | 4 (20.0) |
| Tremor | 4 (19.0%) | 2 (10.0) |
| Abnormal urine | 2 (9.5%) | 2 (10.0) |
| Weight loss | 3 (14.3%) | 1 (5.0) |
| Akathisia | 1 (4.8%) | 2 (10.0) |
| Blurred vision | 1 (4.8%) | 2 (10.0) |
| Dermatologic | 0 (0.0%) | 2 (10.0) |
| Increased salivation | 0 (0.0%) | 2 (10.0) |
| Toxic confusional state | 0 (0.0%) | 2 (10.0) |
| Syncope/dizziness | 0 (0.0%) | 1 (5.0) |
| Tachycardia | 0 (0.0%) | 1 (5.0) |

*Note.* Values are presented as n (%). Each participant was counted once per symptom, regardless of the number of treatment visits at which the symptom was reported. Only side effects reported by at least one participant are shown.

**Supplementary Table 2:** Sensitivity analyses adjusting for medication received (sertraline vs bupropion) and marker assignment (+/- and -/+ vs +/+ vs -/-)

|  |  |  | Primary Model | | Medication-Adjusted | | Marker-Adjusted | |
| --- | --- | --- | --- | --- | --- | --- | --- | --- |
| Analysis | Seed | Region | β | pFDR | β | pFDR | β | pFDR |
| FC ~ change in MADRS | sgACC | R TE1p | 0.029 | 0.023 | 0.030 | 0.014 | 0.029 | 0.104 |
|  | sgACC | R TE1m | 0.030 | 0.023 | 0.032 | 0.014 | 0.027 | 0.104 |
|  | sgACC | L PH | 0.023 | 0.193 | 0.023 | 0.148 | 0.033 | 0.013 |
| FC ~ timepoint (pre vs post) | dACC | L 9p | -1.401 | 0.002 | -1.428 | 0.001 | -1.310 | 0.013 |
|  | dACC | L 8BL | -1.209 | 0.002 | -1.222 | 0.002 | -1.128 | 0.024 |
|  | dACC | L 10d | -1.478 | 0.011 | -1.503 | 0.009 | -1.390 | 0.028 |
|  | dACC | L 47l | -1.154 | 0.012 | -1.182 | 0.010 | -1.158 | 0.028 |
|  | dACC | R 8Av | -1.035 | 0.041 | -1.055 | 0.032 | -0.958 | 0.100 |
|  | dACC | R 8BL | -1.234 | 0.041 | -1.272 | 0.026 | -1.234 | 0.050 |
|  | dACC | R STSda | -1.359 | 0.041 | -1.439 | 0.026 | -1.521 | 0.047 |
|  | dACC | R STSvp | -1.009 | 0.041 | -1.038 | 0.032 | -1.092 | 0.050 |
|  | dACC | R PGi | -1.121 | 0.041 | -1.161 | 0.032 | -1.258 | 0.050 |
|  | dACC | R SFL | -0.792 | 0.082 | -0.807 | 0.062 | -0.898 | 0.050 |
|  | pgACC | R 8BL | -1.688 | 0.005 | -1.716 | 0.004 | -1.581 | 0.014 |
|  | pgACC | R STSda | -1.400 | 0.017 | -1.499 | 0.005 | -1.585 | 0.014 |
|  | pgACC | L PGi | -1.300 | 0.017 | -1.345 | 0.011 | -1.234 | 0.062 |
|  | pgACC | R 47l | -1.363 | 0.020 | -1.377 | 0.018 | -1.262 | 0.068 |
|  | pgACC | L 9p | -1.277 | 0.029 | -1.308 | 0.018 | -1.111 | 0.097 |
|  | pgACC | R 9m | -1.208 | 0.029 | -1.250 | 0.018 | -1.112 | 0.068 |
|  | pgACC | R a32pr | -0.940 | 0.029 | -0.939 | 0.018 | -0.882 | 0.088 |
|  | pgACC | R PGi | -1.216 | 0.029 | -1.282 | 0.018 | -1.179 | 0.068 |
|  | pgACC | L 8BL | -1.140 | 0.031 | -1.148 | 0.021 | -0.914 | 0.134 |
|  | pgACC | L 9m | -1.167 | 0.031 | -1.221 | 0.018 | -1.164 | 0.071 |
|  | pgACC | R STSva | -1.072 | 0.031 | -1.139 | 0.018 | -1.070 | 0.068 |
|  | pgACC | R 8Ad | -0.949 | 0.031 | -0.981 | 0.018 | -0.792 | 0.120 |
|  | pgACC | L 8Ad | -1.048 | 0.034 | -1.125 | 0.018 | -0.860 | 0.120 |
|  | pgACC | L 7m | -1.174 | 0.042 | -1.269 | 0.018 | -1.189 | 0.088 |
|  | pgACC | R 8Av | -0.952 | 0.049 | -0.984 | 0.033 | -0.778 | 0.158 |
|  | pgACC | R STSvp | -0.940 | 0.062 | -0.996 | 0.035 | -0.910 | 0.088 |
|  | pgACC | R PGs | -0.966 | 0.062 | -1.028 | 0.033 | -0.826 | 0.158 |
|  | pgACC | R LO2 | -1.097 | 0.066 | -1.150 | 0.045 | -1.256 | 0.071 |
|  | pgACC | R 7m | -1.036 | 0.066 | -1.101 | 0.043 | -0.920 | 0.158 |

| FC ~ Remission | dACC | R 25 | 2.180 | 0.070 | 2.228 | 0.069 | 2.586 | 0.005 |
| --- | --- | --- | --- | --- | --- | --- | --- | --- |
|  | sgACC | L 8Ad | 2.212 | 0.009 | 2.271 | 0.006 | 2.165 | 0.048 |
|  | sgACC | L TE1m | 2.324 | 0.009 | 2.427 | 0.006 | 2.091 | 0.048 |
| FC ~ Response | pgACC | R TE1m | 3.232 | 0.003 | 3.284 | 0.003 | 2.218 | 0.083 |
|  | pgACC | R TE1p | 2.921 | 0.004 | 2.996 | 0.004 | 2.528 | 0.083 |
|  | pgACC | R PHT | 3.222 | 0.010 | 3.266 | 0.012 | 2.885 | 0.083 |
|  | pgACC | R MIP | 2.513 | 0.058 | 2.598 | 0.047 | 2.160 | 0.263 |
|  | pgACC | R TE2a | 2.079 | 0.058 | 2.137 | 0.047 | 1.406 | 0.263 |
|  | pgACC | R V4t | 2.291 | 0.073 | 2.388 | 0.047 | 2.513 | 0.083 |
|  | pgACC | R PGi | 1.980 | 0.084 | 2.118 | 0.047 | 1.734 | 0.263 |
|  | sgACC | R TE1p | 2.473 | 0.005 | 2.475 | 0.007 | 2.484 | 0.027 |
|  | sgACC | R IFSp | 2.510 | 0.011 | 2.509 | 0.014 | 2.762 | 0.027 |
|  | sgACC | R TE1m | 2.298 | 0.022 | 2.294 | 0.028 | 1.867 | 0.082 |
|  | sgACC | R IFJa | 1.931 | 0.033 | 1.938 | 0.038 | 2.226 | 0.032 |
|  | sgACC | R PHT | 1.871 | 0.037 | 1.875 | 0.038 | 2.170 | 0.044 |
|  | sgACC | R PHA2 | 1.621 | 0.039 | 1.661 | 0.038 | 1.789 | 0.072 |
|  | sgACC | R p47r | 1.828 | 0.045 | 1.878 | 0.040 | 1.818 | 0.089 |
|  | sgACC | R 7Pm | 2.152 | 0.049 | 2.219 | 0.040 | 2.494 | 0.032 |
|  | sgACC | R 47m | 2.106 | 0.049 | 2.130 | 0.046 | 2.426 | 0.080 |
|  | sgACC | R p9.46v | 1.800 | 0.073 | 1.803 | 0.062 | 2.485 | 0.027 |
|  | sgACC | R PGi | 1.560 | 0.073 | 1.677 | 0.046 | 1.376 | 0.156 |

*Note*. Only regions surviving FDR correction (*P*_FDR_ < .05) in at least one model are shown. Medication received and marker assignment (BUP, SERT, Both, or Neither) were examined separately. No significant effects were observed for the hippocampal formation in the primary or sensitivity analyses.
